# Known mechanisms cannot account for at least one third of reduced susceptibility in a diverse collection of non-*aureus* staphylococci

**DOI:** 10.1101/2021.06.22.449369

**Authors:** Heather Felgate, Lisa C. Crossman, Elizabeth Gray, Rebecca Clifford, John Wain, Gemma C. Langridge

## Abstract

**Introduction:** Non-*aureus* staphylococci (NAS) are implicated in many healthcare-acquired infections and an understanding of the genetics of antimicrobial resistance in NAS is important in relation to both clinical intervention and the role of NAS as a reservoir of resistance genes.

**Gap statement:** The burden of antimicrobial resistance in NAS, particularly to clinically relevant antimicrobials, is under recognised.

**Methodology:** We sourced 394 NAS isolates from clinical samples, healthy human volunteers, animals and type cultures and subjected them to agar dilution susceptibility testing against eight antimicrobials. Cefoxitin was used to screen for methicillin resistance in *S. aureus*, as it stimulates expression of *mecA*. We performed whole genome sequencing on 366 isolates and analysed these genotypically for the presence of genetic mechanisms responsible for the phenotypic levels of reduced antimicrobial susceptibility.

**Results:** We observed 175 sequenced isolates with a minimum inhibitory concentration (MIC) of at least 4 μg/ml to cefoxitin, of which 50% (87/175) did not harbour a known *mec* homologue. Eight clinical NAS isolates displayed high daptomycin MICs (>4 μg/ml), with no known mechanism identified. Differences in MICs against erythromycin were attributable to the presence of different resistance genes (*msrA* and *ermC*). In total, 49% (193 /394) of isolates displayed reduced susceptibility to three or more of the antimicrobials tested.

**Conclusions:** The widespread presence of reduced antimicrobial susceptibility in NAS is a concern, with an increased likelihood of (1) harder to treat infections caused directly by NAS, and (2) resistance genes being passed on to other bacteria via horizontal gene transfer, both of which have clinical implications for treatment and management of patients.

## Introduction

The non-*aureus* staphylococci (NAS) represent an important source of nosocomial disease, including prosthetic joint infection (PJI), infective endocarditis and infection in pre-term babies (1). In the UK, over 215,000 joint replacements (hip, knee and shoulder) took place in 2016, with a year-on-year increase of 4% (2). Of these replacements, 1.5% require surgical revision due to infection (2). These infections are most commonly caused by *Staphylococcus* spp., and attributed to NAS in approximately 31% of cases across Europe (3). In our local hospital, the Norfolk and Norwich University Hospital (NNUH), 50% of isolates identified in suspected PJI are NAS.

In clinical microbiology, staphylococci are classified using the coagulase test, with coagulase positive samples overwhelmingly identified as *S. aureus* and coagulase negative samples grouped together under the term coagulase negative staphylococci (CoNS). CoNS is therefore the term found in antimicrobial surveillance data. However, since coagulase negative *S. aureus* strains exist (as do coagulase positive strains of other staphylococcal species), we use the term “non-*aureus* staphylococci” (NAS) to encompass all staphylococci which are not *S. aureus*, regardless of coagulase activity.

There is currently an intense focus upon the presence and spread of bacterial antimicrobial resistance, typified in *S. aureus* by methicillin resistance (MRSA). While the body of literature in antimicrobial resistance research is growing for staphylococci, NAS data remains eclipsed by the focus on *S. aureus*. Few studies have investigated antimicrobial resistance (AMR) in NAS. Those that have suggest 45% of NAS harbour methicillin resistance (4), and that NAS may be resistant to a larger number of antimicrobial classes than *S. aureus* (4, 5).

An understanding of the genetics of AMR in NAS is important in relation to clinical intervention upon diagnosis of PJI (and other infections) as well as understanding the role of NAS as a reservoir of resistance genes. To address this, we assembled a collection of 394 NAS from clinical samples, healthy human volunteers, animals and type cultures and assessed their susceptibility to a range of antimicrobials. We also performed whole genome sequencing to correlate mechanisms of resistance with minimum inhibitory concentrations (MICs).

## Materials & Methods

### NAS collection

Under NHS Research Ethics Committee approval, the Norwich Biorepository banks blood, solid tissue and bacterial isolates from the NNUH and research institutes on the Norwich Research Park, including the University of East Anglia (UEA), and makes these available to the research community. This enabled us to assemble a collection of 380 NAS from a) clinical specimens which were isolated from suspected NAS PJI infections (229, NNUH), b) healthy human volunteers (114, UEA), and c) animal samples (33, UEA) with five having no source recorded. An additional 14 strains of NAS from the National Collection of Type Cultures (NCTC) were supplied by Public Health England.

Isolates were identified using MALDI-TOF (Bruker) to the species level (Table 1). All strains were cultured overnight on TSA plates (Oxoid), checked for contamination and purified. Once purified, the NAS collection was stored as glycerol stocks to be screened for their antimicrobial susceptibility. *Staphylococcus aureus* NCTC 12973 was used as a control.

**Table 1.** Frequency of non-*aureus* staphylococcal species in the study collection

|  |  |
| --- | --- |
| *Staphylococcus auricularis* | 1 |
| *Staphylococcus capitis* | 20 |
| *Staphylococcus caprae* | 2 |
| *Staphylococcus carnosus* | 2 |
| *Staphylococcus chromogenes* | 3 |
| *Staphylococcus cohnii* | 1 |
| *Staphylococcus condimenti* | 1 |
| *Staphylococcus devriesei* | 1 |
| *Staphylococcus epidermidis* | 191 |
| *Staphylococcus equorum* | 1 |
| *Staphylococcus haemolyticus* | 43 |
| *Staphylococcus hominis* | 45 |
| *Staphylococcus jettensis* | 1 |
| *Staphylococcus lugdunensis* | 7 |
| *Staphylococcus massiliensis* | 1 |
| *Staphylococcus microti* | 1 |
| *Staphylococcus muscae* | 1 |
| *Staphylococcus nepalensis* | 1 |
| *Staphylococcus pasteurii* | 4 |
| *Staphylococcus petrasii* | 1 |
| *Staphylococcus pettenkferi* | 1 |
| *Staphylococcus piscifermentans* | 1 |
| *Staphylococcus rostri* | 1 |
| *Staphylococcus saprophyticus* | 22 |
| *Staphylococcus sciuri* | 5 |
| *Staphylococcus simiae* | 1 |
| *Staphylococcus simulans* | 10 |
| *Staphylococcus sp [1]* | 1 |
| *Staphylococcus stepanovicii* | 1 |
| *Staphylococcus succinus* | 1 |
| *Staphylococcus vitulinus* | 3 |
| *Staphylococcus warneri* | 18 |
| *Staphylococcus xylosus* | 1 |

\*Species designated by MALDI-TOF (Bruker)

### Susceptibility testing

To assay the entire NAS collection, five deep well 96-well microplates (VWR) were prepared with 1 ml TSB (Oxoid) per well. Glycerol stocks were used to inoculate the corresponding well. Per plate, one well was designated as a sterility control (TSB only) and one well was inoculated with the *S. aureus* control. After inoculation, plates were sealed and incubated at 37 °C at 180 rpm for a minimum of 10 hours. The experimental design enabled 13.5% of the collection to be tested in duplicate; MIC data was compared and then tabulated (Table S1).

Iso-sensitest agar (Oxoid) was prepared in 250 ml aliquots and autoclaved. Antimicrobial stocks were added to obtain the desired final concentrations once the media had cooled to < 50 °C. For daptomycin, Ca^2+^ was also added at 50μg/ml. The agar antimicrobial mixture was then poured into sterile rectangular plates (Fisher Scientific) and dried.

Per strain, a 1:10 dilution of overnight culture was transferred to a 96 well plate and the OD_600_ was measured. An average OD_600_ was calculated for each column, which was then diluted to approximately OD_600_ 0.6 to generate an inoculum plate for susceptibility testing.

Using a 96-pin multi-point inoculator (Denley), ∼1 μl of inoculum per isolate was stamped onto the agar containing antimicrobials, from the lowest concentration to the highest. Between inoculum plates, the pins were washed in 70 % ethanol for 30 s and allowed to dry before stamping on an antimicrobial-free plate to confirm sterility. Washes were also carried out between antimicrobials using sterile water. All stamped plates were incubated at 37 °C. Isolates found to have reduced susceptibility to daptomycin had their MICs determined for a second time by spotting 10 μl of culture onto TSA plates containing various daptomycin concentrations (supplemented with Ca^2+^ at 50 μg/ml). To increase *mecA* expression, 14 isolates which contained *mecA* but on initial testing showed susceptibility to cefoxitin (MIC < 4 μg/mL) were re-tested on Muller Hinton Agar with 3 % NaCl added alongside a further 6 isolates. Overnight cultures were diluted in PBS and 5 μl spots containing 10^4^ cells were spotted onto plates which were incubated at 35 °C.

Test MIC ranges were determined from British Society for Antimicrobial Chemotherapy (BSAC) surveillance MIC data and EUCAST for coagulase negative staphylococci (CoNS) in 2016 (6). Data for daptomycin were taken from 2010. In μg/ml, the ranges tested were: daptomycin 0.25-2, erythromycin 0.125-256, gentamicin 0.016-64, rifampicin 0.004-0.064, teicoplanin 0.25-16, tetracycline 0.25-256 and vancomycin 1-4. No CoNS surveillance data were available for cefoxitin, hence the test range of 0.25-4 μg/ml was based upon published work (7). Isolates were considered to have reduced susceptibility to the specified antibiotic if they displayed the following MICs: ≥ 4 μg/mL (cefoxitin, teicoplanin, vancomycin); ≥ 2 μg/Ml (tetracycline, erythromycin); ≥ 1 μg/mL (gentamicin, daptomycin); ≥ 0.06 μg/mL (rifampicin).

### Statistical comparison of clinical and non-clinical isolates

Using Prism (GraphPad, San Diego, USA, v 5.04), a Mann-Whitney test was performed (non-parametric test, two-tailed with Gaussian approximation) to compare between the MIC of clinical and non-clinical isolates. Statistical significance was given to a *p* value < 0.05.

### DNA extraction and sequencing

Overnight cultures derived from single colonies were pelleted and resuspended in lysis buffer (Qiagen), transferred to 2 ml lysis matrix B tubes (MPBio) and subjected to bead beating for 15 min at 30 Hz (Tissuelyser II, Qiagen) with RNAse A added. DNA was extracted according to the QiaCube HT protocol with an additional 30 min incubation at 65 °C after proteinase K addition and eluted into Tris-10mM HCL.

Libraries for sequencing were prepared using the Nextera XT DNA Library Prep Protocol and sequenced on the Illumina MiSeq or NextSeq with a loading concentration of 1.8 picomolar.

### Genome analysis

The raw reads were subject to FastQC quality control (8), adapters were trimmed using Trimmomatic [v 0.39] (9) using the supplied NexteraXT adapter sequences. In some cases, read normalisation was performed using BBNorm [v35.85] (10) to remove low coverage contamination. The lowest coverage cutoff level parameter used was dependent on the total coverage of the sequence since some sequencing runs had a high difference in coverage level across the run. Finally, reads were concatenated if they originated from Illumina NextSeq since this platform produces eight reads per sample, four forward and four reverse. A total of 364 samples passed QC and were suitable for downstream analysis. These reads were used as described below. Sequences are available from the European Nucleotide Archive, under project PRJEB31403.

To determine which antimicrobial resistance genes and associated individual mutations were present in each of our 364 NAS genomes, reference gene sequences were downloaded from CARD v2.0.0 (11) and used as input to ARIBA v2.13.2 (12) which generates local assemblies from sequence reads and reports back which reference genes (and individual mutations) are identified, with a minimum percent identity cut off at 90% (Table S5). For genes where ‘partial’ or ‘interrupted’ was reported, this was not considered sufficient evidence for intact gene presence. The tabulated results were evaluated for gene and mutation presence/absence relative to MIC per antimicrobial. Twelve NCTC sequences were downloaded as genome assemblies from the European Nucleotide Archive (accessions: SAMEA4364213; SAMEA4364214; SAMEA4384234; SAMEA4384058; SAMEA4384237; SAMEA4384064; SAMEA4412661; SAMEA4384059; SAMEA4384235; SAMEA4384060; SAMEA4384339; SAMEA4384403) and analysed by ABRIcate v0.9.7 (13) using CARD v2.0.0 (11) as the reference database with a minimum DNA coverage of 90%. NCTC 13831 and 13837 sequence data was not available at time of sequencing and therefore these isolates were sequenced as described above for the main NAS collection. Protein level conservation was assessed using BLAST v2.10.1 against the NCBI AMR database. Hits were recorded for greater than 40% identity at the protein level over 80% of the query and subject sequence.

## Results & Discussion

Our NAS collection comprised over 30 species of *Staphyloccocus*, including at least 10 isolates of *S. epidermidis, S. capitis, S. haemolyticus, S. hominis, S. saprophyticus, S. simulans* and *S. warneri* (Table 1). Isolates were collected over a four-year period from 2013 to 2016. The range of antimicrobials tested were selected based upon clinical relevance and availability (Table S1). Observed MIC distributions per antimicrobial are shown in Figure 1.

**Figure 1.**
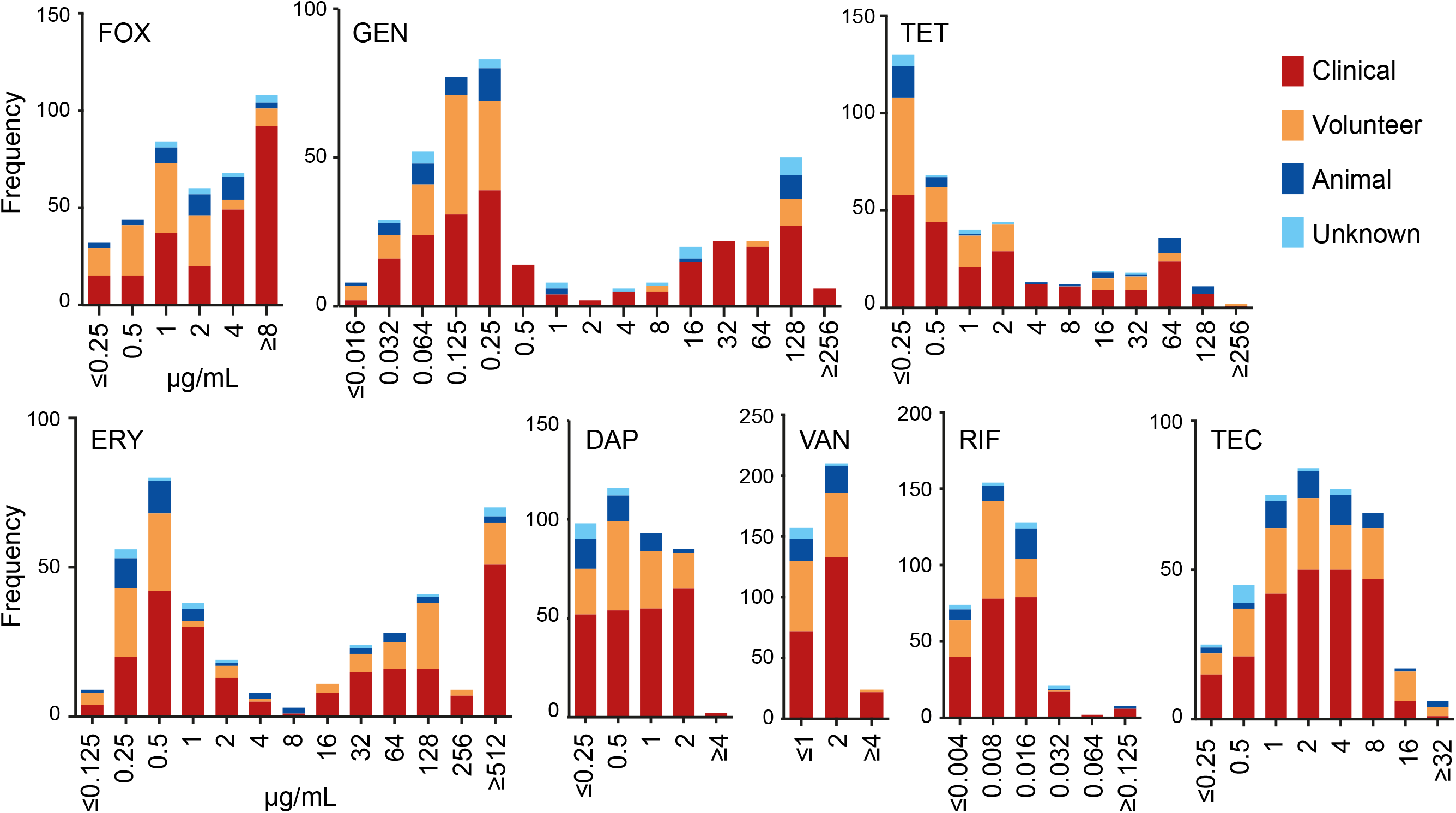
Antibiotic susceptibility of NAS collection MIC distributions for 394 NAS isolates grown in the presence of antimicrobials. FOX cefoxitin, GEN gentamicin, TET tetracycline, ERY erythromycin, DAP daptomycin, VAN vancomycin, RIF rifampicin and TEC teicoplanin.

### *Cefoxitin screening does not correlate with* mecA *presence in clinically relevant NAS*

Cefoxitin is used to screen for methicillin resistance in *S. aureus* as it induces *mecA* expression. However, while methicillin resistant *S. aureus* (MRSA) have a high public profile, much less is known about methicillin resistant NAS (MRNAS). EUCAST guidelines state that for MRSA “cefoxitin is a very sensitive and specific marker of *mecA/mecC*-mediated methicillin resistance including in heterogeneous expressing strains and is the agent of choice” (14). In this collection, we found 194/394 (49%) displayed reduced susceptibility to cefoxitin with MICs ≥ 4 μg/ml (Table S1). The vast majority of these isolates were from clinical samples (FOX Figure 1, Table S1) but analysis at the nucleotide level (Table S2 and S3) indicated that only 88 out of 175 (50 %) sequenced isolates with an MIC ≥ 4μg/ml harboured a known *mecA*. MecA is extremely well-conserved and searching at the protein level yielded the same results (Table S4). Other *mec* elements were also identified (e.g. *mecC, mecI* and *mecR1*) but only ever in addition to *mecA*. Breaking this down by species, 20/21 *S. saprophyticus* isolates with cefoxitin MIC ≥ 4μg/ml harboured no *mecA* (Figure 2). No *mecA* was detected in eleven species with high cefoxitin MICs and for *S. hominis, S. warneri* and *S. haemolyticus*, the percentage of the population that exhibited MIC ≥ 4μg/ml with no *mecA* was between 28 and 67% (Figure 2). Our results support cefoxitin as a good indicator of *mecA* presence in *S. epidermidis* (7), but suggest that it performs poorly in the less common, but still clinically relevant NAS. In addition, we observed 14 cases where the presence of *mecA* did not result in an MIC ≥ 4μg/ml. These were re-tested (alongside 6 others) under conditions designed to encourage *mecA* expression and resulted in increased MICs in all isolates, however 7/20 remained <4 ug/mL (Table S1).

**Figure 2.**
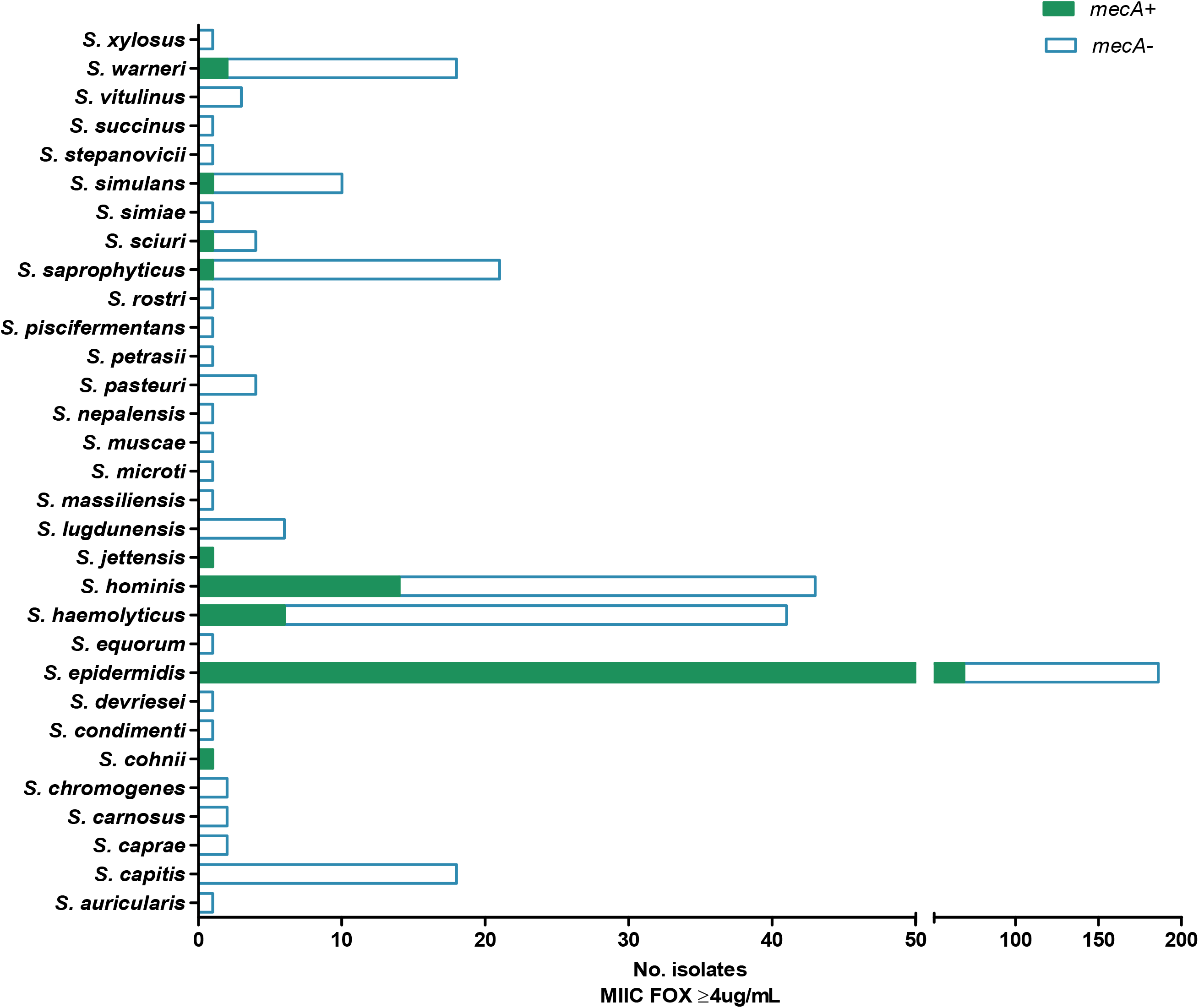
Presence of *mecA* in relation to high cefoxitin MIC. Per staphylococcal species, bars display the total number of sequenced isolates found to have a cefoxitin MIC of ≥4 μg/ml, in relation to the presence (solid green) and absence (blue outline) of *mecA*.

### Reduced susceptibility only partly explained by known mechanisms

When exposed to gentamicin and tetracycline, isolates could broadly be divided into two populations, displaying susceptible or reduced susceptibility phenotypes (Figure 1 GEN and TET). In isolates displaying an MIC ≥1 μg/ml for gentamicin, 49/130 isolates harboured *aac(6’)-le-aph(2”)-la* (Table 2) which is associated with gentamicin resistance in *Enterococcus* (15, 16) but has also been observed in *Staphylococcus* (17, 18). A total of 12 isolates had a match for *aph(3’)IIIa*, but only five of them were associated with reduced susceptibility.

**Table 2.** Genetic mechanisms identified using ARIBA/ABRIcate and the CARD database compared to the MIC data (of sequenced isolates only, partial and interrupted sequences are not included, see Table S2)

| *Antimicrobial* | *Mechanism*<br>*(Accession*<br>*No.)* | *No. isolates above breakpoint*<br>*MIC ($\geq 2 \mu\text{g/ml}$)* |  | *No. isolates below*<br>*breakpoint* |
| --- | --- | --- | --- | --- |
| TET | *tetK*<br>(NC_013452) | 48/148 |  | 5/223 |
|  | *tetL*<br>(M11036.0) | 1/148 |  | 0/223 |
|  | *tetM*<br>(AM180355) | 1/148 |  | 0/223 |
|  | **All\*** | **50/148 (33.8%)** |  | **5/223 (2.2%)** |
|  |  | *No. isolates above breakpoint*<br>*MIC ($\geq 1 \mu\text{g/ml}$)* |  |  |
| GEN | *aac(6')-Ie-aph(2'')-Ia*<br>(NC_005024) | 49/130 |  | 6/246 |
|  | *aph(3)IIIa*<br>(CP004067) | 5/130 |  | 7/246 |
|  | **All\*** | **49/130 (37.7 %)** |  | **12/246 (4.9 %)** |
|  |  | *MIC*<br>*$\geq 512 \mu\text{g/ml}$* | *MIC*<br>*$\geq 2 - 256 \mu\text{g/ml}$* |  |
| ERY | *msrA*<br>(NC_022598.1) | 6/66 | 75/135 | 9/173 |
|  | *ermC*<br>(M12730) | 42/66 | 13/135 | 9/173 |
|  | *ermA*<br>(NC_009632) | 6/66 | 1/135 | 0/173 |
|  | *emeA*<br>(AB091338) | 1/66 | 0/135 | 0/173 |
|  | **All\*** | **46/66 (69.7 %)** | **88/135 (65.2 %)** | **18/173 (10.4 %)** |

\*Some isolates harboured multiple resistance genes. TET tetracycline, GEN gentamicin, ERY erythromycin.

Six isolates that contained *aac(6’)-le-aph(2”)-la* displayed susceptible MICs, making them the equivalent of major errors (MEs) in public health terms, as the isolates were genotypically resistant but phenotypically susceptible (19). Accordingly, the 81/130 isolates with reduced susceptibility (≥1 μg/ml) that harboured no *aac(6’)-le-aph(2”)-la* represented the equivalent of very major errors (VMEs) as they were genotypically susceptible but phenotypically resistant (19). This is highly suggestive of novel mechanisms of resistance and was a feature of other antimicrobials tested (Figure 3).

**Figure 3.**
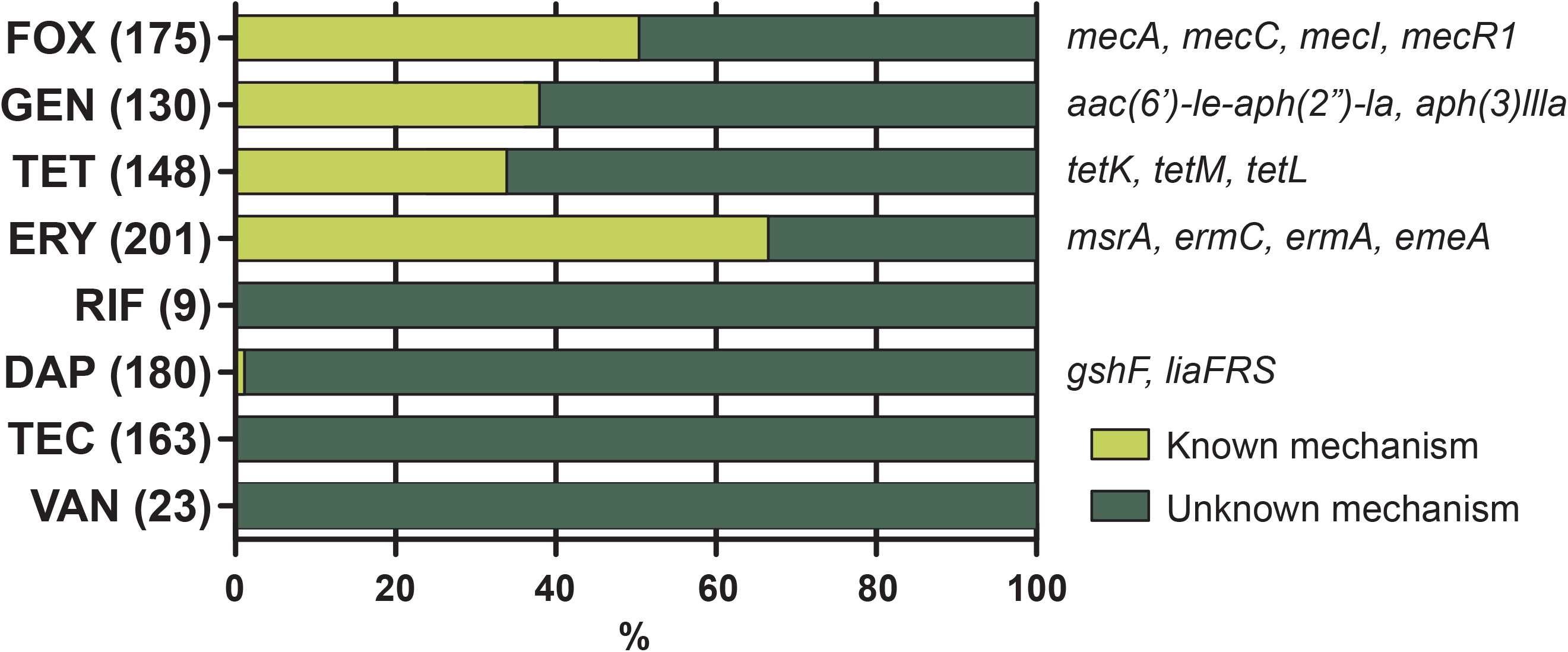
Mechanisms of resistance Percentage of known genetic mechanisms identified in sequenced NAS isolates with reduced susceptibility. Number of isolates with reduced susceptibility per antimicrobial given in parentheses. Known mechanisms found per antimicrobial are shown in italics. FOX cefoxitin, GEN gentamicin, TET tetracycline, ERY erythromycin, DAP daptomycin, VAN vancomycin, RIF rifampicin and TEC teicoplanin.

To identify whether efflux pumps might play a role these phenotypes, we assessed the ARIBA output for the staphylococcal-specific *norABC, mgrA, mepR* and *qac* genes (20). In the sequenced NAS collection 168/378 (44.4 %) contained *norA*, however of these less than two thirds (75/130) had reduced susceptibility to gentamicin.

According to the Comprehensive Antimicrobial Resistance Database [CARD] (11), *tetK* is by far the most common tetracycline resistance mechanism in *S. aureus* and *S. epidermidis* (10-20%), followed by *tetL* (<1%) and *tetM* (<1%). This was borne out in our NAS collection, where 48/148 (32.4 %) isolates with MICs ≥ 2 μg/ml of tetracycline contained *tetK*, as compared to 5/223 (2.2 %) with MICs below 2 μg/ml. One animal isolate with an MIC of 16 μg/ml carried *tetL* and one clinical isolate with an MIC of 64 μg/ml carried *tetM*; neither had any other tetracycline resistance genes. Again, this demonstrated that 98/148 isolates displayed a reduced susceptibility phenotype that did not associate with a known resistance determinant, indicative of uncharacterised resistance mechanisms.

The distribution of erythromycin phenotypes was more complex. With this antimicrobial, we observed both susceptible isolates and those with reduced susceptibility, but the latter appeared to consist of two populations, one with MICs between ≥2-256 μg/ml and one with MICs ≥512 μg/ml (Figure 1 ERY). We had sequence data available from 135 of the ≥2-256 μg/ml population and 66 of the ≥512 μg/ml population, and identified the presence of a resistance gene (*ermA, ermC, msrA)* in 65.9 % (89/135) of the ≥2-256 μg/ml population and 55/66 (83.3 %) of the ≥512 μg/ml population (Table 2). Our results indicated that the presence of *ermC* rather than *msrA* was the major cause of MICs exceeding 256 μg/ml. Although rare (n=4), harbouring both genes resulted in an MIC ≥ 512 μg/ml in three cases and 128 μg/ml in the other. In isolates with an MIC of ≥ 512 μg/ml, *qac* was identified 36 times. In 23 of these cases *ermC* was also present; *qac* was only found twice with no other known erythromycin resistance mechanisms present. A total of 101 isolates with an MIC ≥2 μg/ did not contain *qac*.

For daptomycin, approximately half the collection displayed reduced susceptibility (MIC ≥ 1 μg/ml, Figure 1 DAP and Table S1). A small subset, comprising eight isolates from clinical samples only, displayed MICs ≥ 4 μg/ml; such high MICs to daptomycin have not been previously reported, according to the European Committee on Antimicrobial Susceptibility Testing (EUCAST) and The British Society for Antimicrobial Chemotherapy (BSAC) surveillance data. These MICs were repeated a second time and confirmed. This is concerning given that daptomycin is a current therapeutic choice for treating soft tissue infections caused by NAS (21). Seven of these isolates were sequenced and our ARIBA analysis (Table 3) indicated only a single *S. epidermidis* isolate contained genes implicated in daptomycin resistance: *gshF* and *liaFRS* with an MIC of 1 μg/ml, the remaining 169 isolates with an ≥1 μg/ml MIC did not harbour any of these genes. Several mutations or genes are associated with daptomycin resistance in *S. aureus* (including *mprF*, and SNPs in *rpoC*) but none of these were identified in the NAS collection (6, 22). More recently, SNPs in *walK* have also been associated with daptomycin resistance in *S. aureus* and *S. epidermidis* (23). The three *S. aureus* SNPs are present in CARD and were not identified in our collection. The V500F mutation from *S. epidermidis* (23) was also not identified in our *S. epidermidis* with DAP MICs ≥ 4 μg/ml. Whilst *walK* was identified by protein BLAST as present across the NAS collection (as expected for an essential gene (24)), sequence variation was observed at the protein level which prevents SNPs observed in *S. aureus* or *S. epidermidis* being extrapolated to NAS. We therefore conclude that there are potentially novel daptomycin resistance mechanisms present in these strains.

### Resistance to vancomycin found in clinical samples

Vancomycin is a treatment option in prosthetic joint infection, and 94 % of isolates had an MIC below 4 μg/ml (Figure 1 VAN). However, of the 24 isolates with reduced susceptibility, 22 (92 %) came from clinical samples and only 2/24 were found in healthy volunteers. This is indicative of a wider trend, where isolates associated with clinical samples had significantly higher MICs (p < 0.005) than non-clinical isolates for cefoxitin, erythromycin, gentamicin, tetracycline, daptomycin and vancomycin (Figure S1). Given the importance of NAS in nosocomial infections, this is a worrying prospect both in terms of what is presenting in the clinic and also the possibility of AMR gene transfer into organisms more capable of causing infection, including *S. aureus*. In addition, no known mechanisms of resistance were identified for vancomycin, rifampicin or teicoplanin (Figure 3 and Table S3).

### Over half of the NAS collection displayed susceptibility to multiple antimicrobials

Out of the all the isolates tested, 48 % of (192/394) had reduced susceptibility to three or more antimicrobials. Twenty-five isolates had reduced susceptibility to six antimicrobials, and three isolates had reduced susceptibility to seven antimicrobials; of these 24/25 and 3/3 were isolated from clinical samples (Table S1). The implications of these are difficult to treat infections and potentially a large reservoir of staphylococcal resistance genes within the patient under antimicrobial treatment.

### Animal isolates have similar MIC distributions to human isolates

It is generally acknowledged that the presence of reduced susceptibility in microorganisms isolated from animals can impact upon public health if those organisms also cause infection in humans (25, 26). In our collection there were 40 NAS isolated from animals (7 NCTC strains), of which we obtained genome sequences from 23. Although in much fewer numbers than the human isolates in the collection, the animal isolates displayed very similar MIC distributions and harboured corresponding genetic mechanisms. This does not rule out the possibility that animals could be a reservoir of AMR for staphylococci.

## Conclusion

Genome analysis of isolates displaying MICs to cefoxitin of ≥ 4 μg/ml indicated that approximately half harboured the *mecA* element. The absence of *mecA* from the other half suggests that other mechanisms are likely present. This was apparent across many of the antimicrobials tested as between 0 and 65 % of phenotypic resistance in clinical isolates could be attributed to known resistance mechanisms. The remaining 35-100 % suggests that there are potentially numerous unknown mechanisms underpinning NAS resistance, which warrant further investigation.

## Acknowledgements

We thank David Livermore and Iain McNamara for productive discussions on antimicrobial usage.

## Transparency declarations

None to declare.

## Supplementary data

Table S1. MIC characterisation of the NAS collection

MIC data for all isolates tested; all MIC are shown in μg/ml; FOX cefoxitin, GEN gentamicin, TET tetracycline, ERY erythromycin, DAP daptomycin, VAN vancomycin, RIF rifampicin and TEC teicoplanin.

Table S2. ARIBA characterisation of all sequenced NAS isolates, grouped by relevant antimicrobial.

Table S3. Summary of ARIBA/ABRIcate match data for all sequenced isolates.

Table S4. Comparison of gene presence (using ARIBA) and protein BLAST presence. Data includes all genes identified in the CARD database.

Table S5. Full list of all reference genes/mutations from the CARD database analysed for presence/absence.

Figure S1. MICs in clinical and non-clinical isolates

Box and whisker plot of MIC distribution per antimicrobial for clinical (orange, CS) and non-clinical (yellow, NC) isolates. Thick black bar indicates the median MIC which resided within the Interquartile Range. The extreme lines are represented as dotted lines and indicate the data outside the upper (75%) and lower (25%) quartiles, open circles represent potential outliers. Levels of significance from Mann-Whitney U test denoted by **** (p<0.0001), *** (p<0.005), ** (p=0.001) and * (p=0.01).

