## Supplementary figures and images for "Known mechanisms cannot account for at least one third of reduced susceptibility in a diverse collection of non-*aureus* staphylococci"

### Figure S1

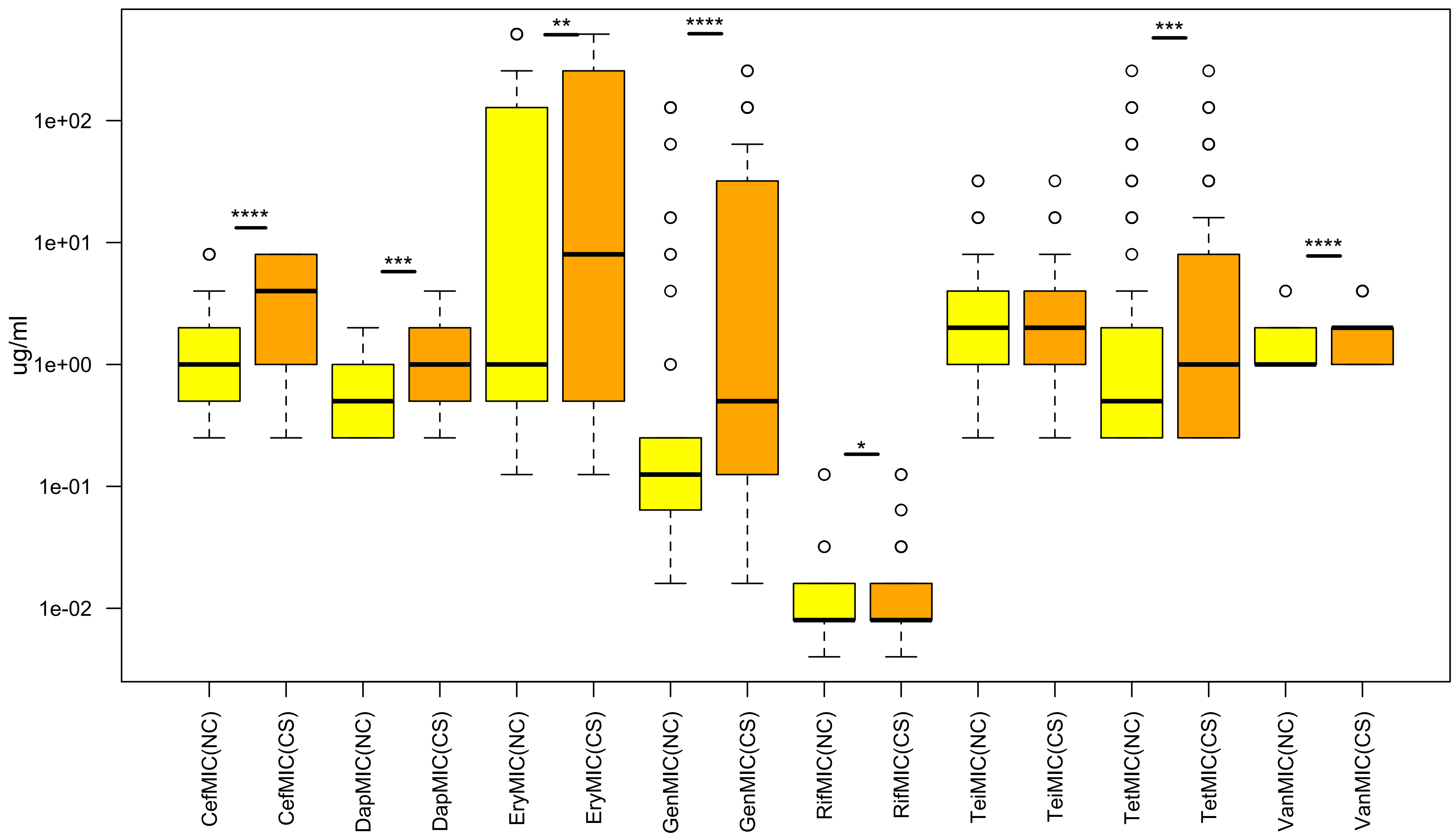
